# NPF4.6-mediated ABA transport modulates stomatal responses to fluctuating light

**DOI:** 10.64898/2025.12.30.697139

**Authors:** Naoya Katsuhama, Ryo Yamauchi, Yumiko Takebayashi, Masami Yokota Hirai, Mitsunori Seo, Wataru Yamori

**Affiliations:** Graduate School of Agricultural and Life Sciences, The University of Tokyo, Nishitokyo, Tokyo, Japan; RIKEN Center for Sustainable Resource Science, Yokohama, Kanagawa, Japan; Tropical Biosphere Research Center, University of the Ryukyus, Nishihara, Okinawa, Japan

**Keywords:** *Arabidopsis thaliana*, NPF4.6, ABA transport, hormone signaling, fluctuating light, photosynthetic induction, stomatal conductance

## Abstract

Stomatal responses to fluctuating light (FL) strongly influence photosynthetic efficiency and biomass accumulation in seed plants. The abscisic acid (ABA) transporter NPF4.6 (also known as NRT1.2 or AIT1) mediates hormone import into guard cells, but its physiological roles in leaves remain unclear. Here, we examined the functions of NPF4.6 in *Arabidopsis thaliana* by characterizing two independent loss-of-function mutants (*npf4.6*) under both steady and FL conditions. Compared with wild type, *npf4.6* showed a 21–43% reduction in the half-time of stomatal opening and a 26–32% increase in the mean net CO_2_ assimilation rate under simulated field light conditions, resulting in a 41–48% increase in shoot biomass. In contrast, no marked differences in photosynthetic efficiency were observed between the wild type and *npf4.6* under drought stress, suggesting that the role of NPF4.6 is more pronounced under well-watered, dynamic-light conditions. Overall, these findings demonstrate that NPF4.6 contributes to the fine-tuning of stomatal kinetics under variable light environments, and highlight hormone transport as a promising target for improving photosynthesis and growth in the field.

## Introduction

Plants in natural environments experience rapid fluctuations in light caused by clouds, canopy movement, and sunflecks (Pearcy, 1990; Slattery et al., 2018). Efficient use of fluctuating light (FL) requires rapid adjustment of both stomatal and biochemical components of photosynthesis (Yamori, 2016). However, a major limitation under FL is the slow stomatal opening, which delays CO_2_ diffusion into the mesophyll and reduces photosynthetic induction (Kaiser et al., 2015; Lawson and Matthews, 2020; Yamori et al., 2020; Tanigawa et al., 2024; Yoshiyama et al., 2024). Thus, it is well established that faster photosynthetic induction, particularly through improved stomatal responses to light, can enhance carbon gain and biomass accumulation in several plant species, including crops (Acevedo-Siaca et al., 2020; Kimura et al., 2020; Ermakova et al., 2023).

Stomata are pores located in the epidermis that regulate CO_2_ uptake and water loss through complex interactions with environmental and hormonal signals (Leakey et al., 2019; Lawson and Matthews, 2020). Among phytohormones, abscisic acid (ABA) is a key regulator of stomatal closure during drought (Kuromori et al., 2018; Hsu et al., 2021). While ABA’s role during soil water deficit is well established, recent studies have shown that ABA also modulates stomatal behavior in response to fluctuations in light, CO_2_ concentration, and air vapor pressure deficit, independently of soil water status (Merilo et al., 2018; Shimadzu et al., 2019; Hsu et al., 2021; Yang et al., 2024). For example, although ABA signaling is not necessary for dark-induced stomatal closure, ABA receptor mutants (*q1124* and *s112458*) and the biosynthesis mutant *nced3/5* show delayed responses to darkness (Pridgeon and Hetherington, 2021). In addition, in the ABA-deficient mutant *aba3-1*, stomata fail to close under low relative humidity, but this phenotype is rescued by guard cell-specific expression of *ABA3*, indicating that guard cells autonomously produce ABA rather than responding to root-derived drought stress (Bauer et al., 2013). Together, these findings suggest that ABA not only functions as a binary signal for stomatal closure under drought, but also modulates the dynamics of stomatal responses under rapidly changing environmental conditions.

The effects of ABA depend not only on its synthesis and catabolism but also on its spatial distribution within the plant, which is regulated by specific transporters (Kuromori et al., 2018; Morales de Los Ríos et al., 2025). ABA is synthesized in multiple tissues, including roots, leaf vascular tissues, and guard cells, with vascular tissues likely serving as major sites of its production (Kuromori et al., 2018). Because guard cells are located at the leaf surface and ABA is mainly produced in internal vascular tissues, intercellular transport is required to deliver ABA to guard cells and regulate stomatal behavior (Kuromori et al., 2018). Several classes of ABA transporters have been identified, including the ABC transporter family (e.g., AtABCG25 and ABCG40) and the NITRATE TRANSPORTER 1/PEPTIDE TRANSPORTER FAMILY (NPF) (Chiba et al., 2015; Kuromori et al., 2018; Morales de Los Ríos et al., 2025).

NPF4.6 (also known as NRT1.2 or AIT1) was first identified as a low-affinity nitrate transporter in *Arabidopsis thaliana* roots (Huang et al., 1999) and later shown to act as an ABA importer that regulates stomatal aperture by mediating ABA uptake into guard cells (Kanno et al., 2012; Shimizu et al., 2021). Notably, *NPF4.6* is expressed in guard cells, and its guard cell-specific complementation partially rescued the reduced leaf temperature of the *aao3 npf4.6* double mutant, indicating that NPF4.6 functions within guard cells to promote ABA-dependent stomatal closure (Shimizu et al., 2021). Recent work has shown that *abcd25* and *abcg40* mutants exhibit faster stomatal opening upon irradiation, possibly due to reduced ABA transport to guard cells, thereby lowering guard-cell ABA levels (Shimadzu et al., 2019). Given that NPF4.6 mediates ABA uptake into guard cells, it may similarly regulate stomatal aperture under FL. Specifically, maintenance of guard-cell ABA by NPF4.6 could restrict stomatal opening under increasing light, whereas loss of NPF4.6 function might accelerate stomatal opening and enhance CO_2_ uptake during photosynthetic induction. However, the impact of NPF4.6 on gas exchange in leaves under realistic light conditions remains untested.

Here, we examined the contribution of NPF4.6 to stomatal regulation under fluctuating light, focusing on gas-exchange dynamics in photosynthetic tissues. Understanding the function of NPF4.6 in this context provides insights into ABA transport during environmental acclimation and its potential contribution to carbon gain in field-grown crops.

## Results

### Accelerated photosynthetic induction in npf4.6 mutants

To assess the effect of NPF4.6-mediated ABA transport on stomatal opening and photosynthetic induction, we compared the light response of wild type (WT) and *npf4.6* mutants (*npf4.6-1* and *npf*4.6-*2*) (Fig. 1). The steady-state net CO_2_ assimilation rate (*A*), leaf chlorophyll content, and Rubisco content did not differ significantly between the genotypes tested (Fig. 1C, Fig. 2I, J). However, *npf4.6* mutants tended to show higher steady-state stomatal conductance (*g*_sw_) (Fig. 1G, Fig. 2G) and significantly higher intercellular CO_2_ concentration (*C*_i_) (Fig. 1I, K, Fig. 2H). Also, intrinsic water use efficiency (*iWUE*) was significantly lower in *npf4.6* than in WT (Fig. 1J, L), reflecting a smaller increase in *A* relative to *g*_sw_.

**Figure 1.**
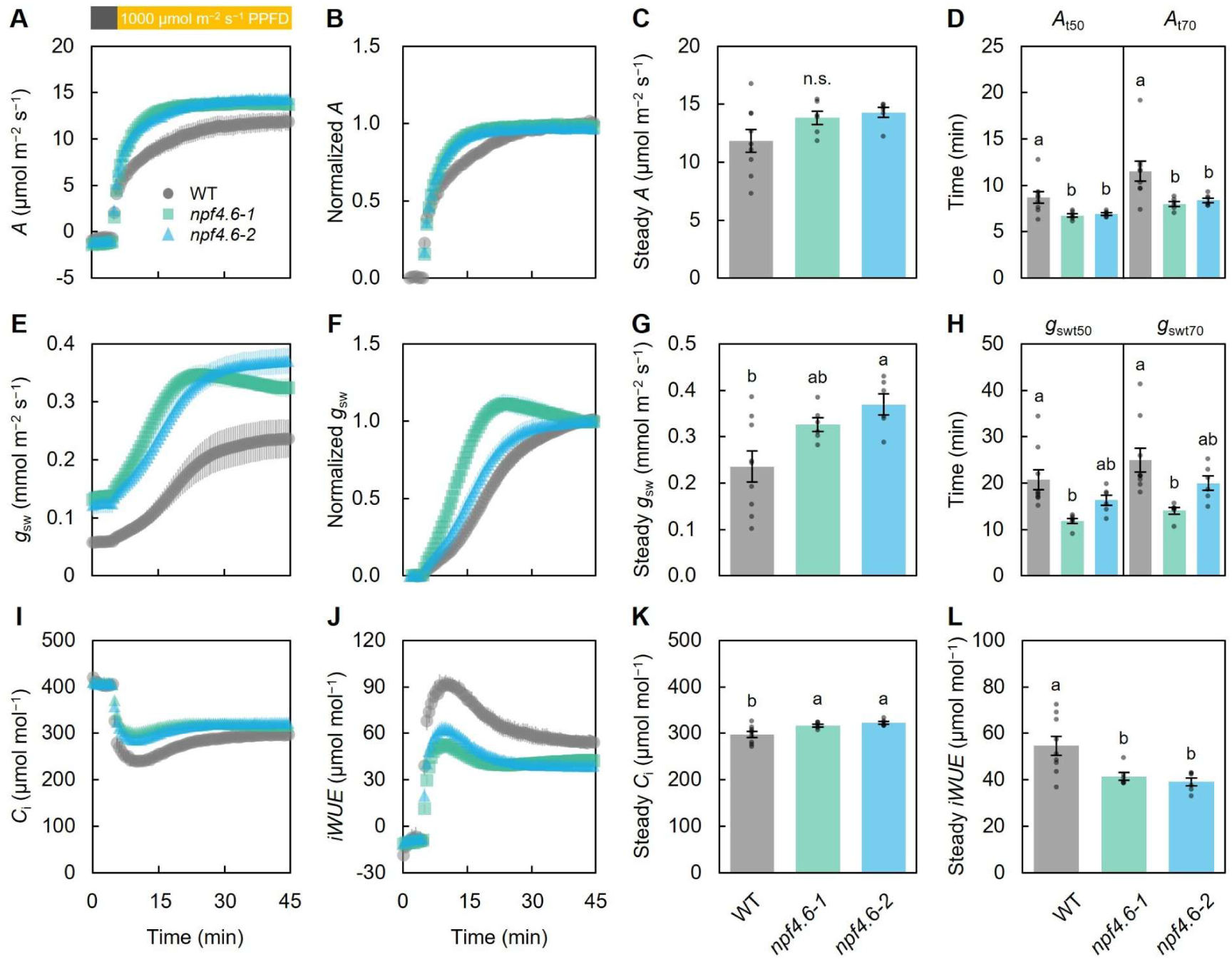
Photosynthetic induction under a step increase in light intensity. (A–D) Net CO_2_ assimilation rate (*A*), (E–F) stomatal conductance (*g*_sw_), (I, K) intercellular CO_2_ concentration (*C*_i_), and (J, L) intrinsic water use efficiency (*iWUE*) of wild type (WT) and *npf4.6* mutants during a shift from dark to high light (PPFD of 1000 μmol m^−2^ s^−1^). (B, F) Normalized *A* and *g*_sw_. (C, G, K, L) Steady-state *A*, *g*_sw_, *C*_i_, and *iWUE* at 35–40 min after light onset. (D, H) Time to reach 50% (_t50_) and 70% (_t70_) of maximum *A* and *g*_sw_. Values are mean ± SEM. Different letters indicate significant differences at *P* < 0.05 by Tukey-Kramer HSD test (*n* = 9 for WT, *n* = 6 for *npf4.6* mutants).

**Figure 2.**
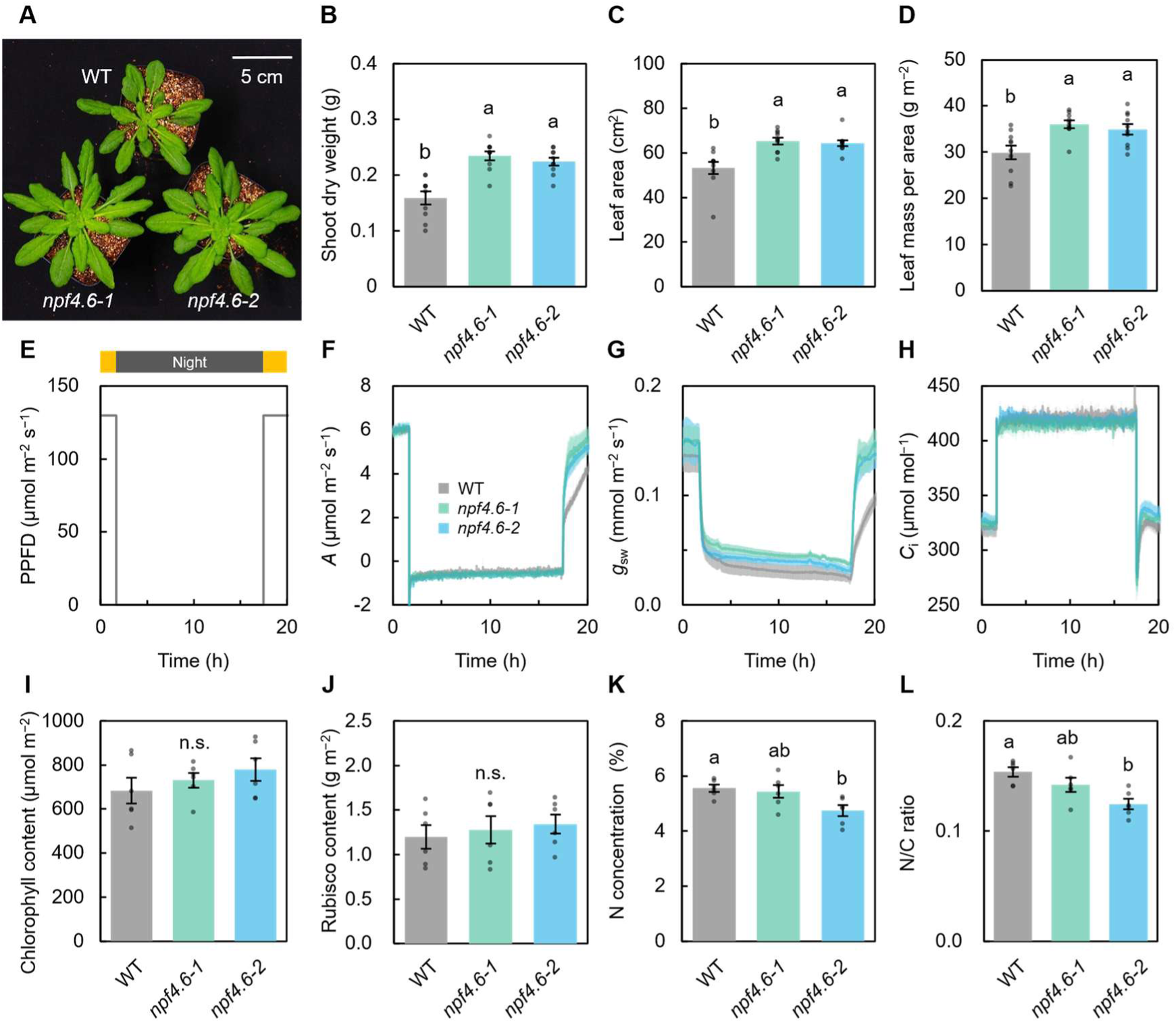
Plant growth and diurnal photosynthesis under short-day conditions with constant light. (A) Representative growth at 50 days after planting. Scale bar = 5 cm. (B–D) Shoot dry weight, projected leaf area, and leaf mass per area. (E) Light pattern used for measurement. (F–H) Diurnal changes in *A*, *g*_sw_, and *C*_i_. (I–L) Total chlorophyll content, Rubisco content, N concentration, and N/C ratio in mature leaves. Values are mean ± SEM. Different letters indicate a significant difference at *P* < 0.05 by Tukey-Kramer HSD test (*n* = 10 for A–D, *n* = 4 for E–H, *n* = 6 for I–L).

When normalized to steady-state values, both *A* and *g*_sw_ increased more rapidly in *npf4.6* than in WT (Fig. 1B, F). The times required to reach 50% and 70% of the steady-state *A* (*A*_t50_ and *A*_t70_) were significantly shorter in *npf4.6* (Fig. 1D). Similarly, the times required to reach 50% and 70% of the steady-state *g*_sw_ (*g*_swt50_ and *g*_swt70_) were marginally shorter in *npf4.6* (Fig. 1H). Together, these results indicate that *npf4.6* mutants show faster stomatal opening and photosynthetic induction in response to light transitions.

### Enhanced photosynthesis in npf4.6 mutants under natural FL

To test whether accelerated photosynthetic induction in *npf4.6* mutants improves dynamic photosynthesis under natural FL, we monitored leaf gas exchange under 11 h fluctuating light patterns mimicking middle-canopy light fluctuations in paddy fields (Fig. 3). Under FL, *npf4.6* mutants showed significantly higher mean *A* and a marginally higher (*P* = 0.057–0.074) *g*_sw_ compared with WT (Fig. 3B, C). More specifically, *npf4.6* mutants maintained relatively higher *A* and *g*_sw_ in the afternoon (Fig. 3E, F). Furthermore, *iWUE* was lower in *npf4.6* mutants (Fig. 3D). These results indicate that faster photosynthetic induction enhances the photosynthetic response to dynamic light, resulting in higher photosynthesis throughout the day.

**Figure 3.**
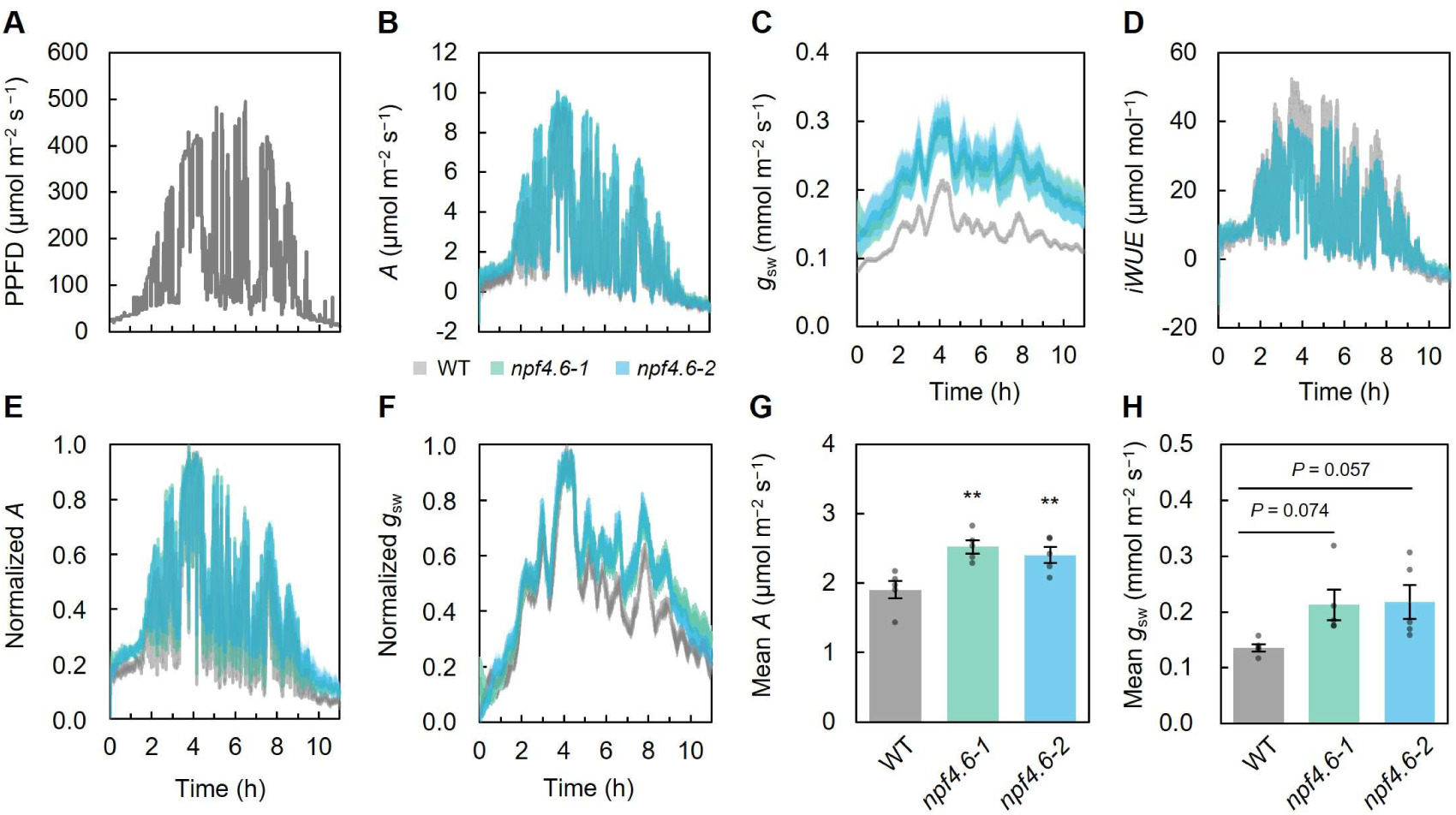
Dynamic photosynthesis of WT and *npf4.6* mutants under simulated field light conditions. (A) Field-measured light fluctuations were scaled to a maximum PPFD of 500 μmol m^−2^ s^−1^ and shortened to 11 h for measurement. (B–D) *A*, *g*_sw_, and *iWUE* were measured at 10-s intervals. (E) Normalized *A* and *g*_sw_. (G, H) Mean *A* and *g*_sw_ across the photoperiod. Values are mean ± SEM. Significant differences from WT by Dunnett’s test are indicated by ** for *P* < 0.01 and * for *P* < 0.05 (*n* = 5).

### Growth promotion in npf4.6 mutants

To examine whether the accelerated stomatal response and photosynthetic induction in *npf4.6* enhances plant growth, plants were grown under a short-day condition with constant light at PPFD of 130 μmol m^−2^ s^−1^ (Fig. 2). *npf4.6* mutants produced significantly greater shoot biomass than WT (Fig. 2A, B), supported by increased projected leaf area (Fig. 2C). Leaf mass per area (LMA) was also significantly higher in *npf4.6* mutants (Fig. 2D). Root dry weight was significantly greater as well (Fig. S1), suggesting that greater carbon fixation and root uptake contributed to whole-plant growth. Although one *npf4.6-2* showed significantly reduced leaf N concentration and N/C ratio, this trend was not consistent among the mutants (Fig. 2K, L).

We also measured diurnal photosynthetic responses under the growth chamber light regime (Fig. 2E). While gas exchange was comparable in the dark, *npf4.6* mutants showed higher *g*_sw_ and *C*_i_ during the first hours of the light period, resulting in higher *A* (Fig. 2F–H). However, no difference in steady-state *A* was found at the end of the day (Fig. 2F). These results suggest that enhanced photosynthetic induction at light onset improved carbon assimilation and contributed to greater growth in *npf4.6* mutants under short-day conditions.

### ABA transport function of NPF4.6

To further examine the role of NPF4.6 in ABA transport and stomatal regulation, we analyzed stomatal closure of 4- to 5-week-old plants in response to exogenous ABA using thermal imaging (Fig. 4). In the absence of ABA, leaf temperatures were comparable between WT and *npf4.6* mutants (Fig. 4B), consistent with earlier reports (Kanno et al., 2012; Shimizu et al., 2021). However, after ABA application through the roots and overnight incubation, *npf4.6* mutants showed lower leaf temperatures than WT, indicating less ABA-induced stomatal closure (Fig. 4A, B). Previous studies have shown that NPF4.6 mediates ABA uptake into guard cells, but it is also expressed in leaf and root vascular tissues (Kanno et al., 2012; Shimizu et al., 2021). Therefore, altered stomatal response in *npf4.6* mutants to the root-applied ABA could result from impaired long-distance ABA transport from roots to leaves via vascular tissues. To test this, we quantified ABA contents in xylem sap following root ABA treatment in 7- to 8-week-old plants (Fig. 4C). However, no significant differences in xylem sap ABA concentration were found between WT and *npf4.6* mutants (Fig. 4C), suggesting that reduced stomatal responsiveness in the mutants reflects impaired ABA uptake into guard cells rather than defects in root-to-shoot systemic transport.

**Figure 4.**
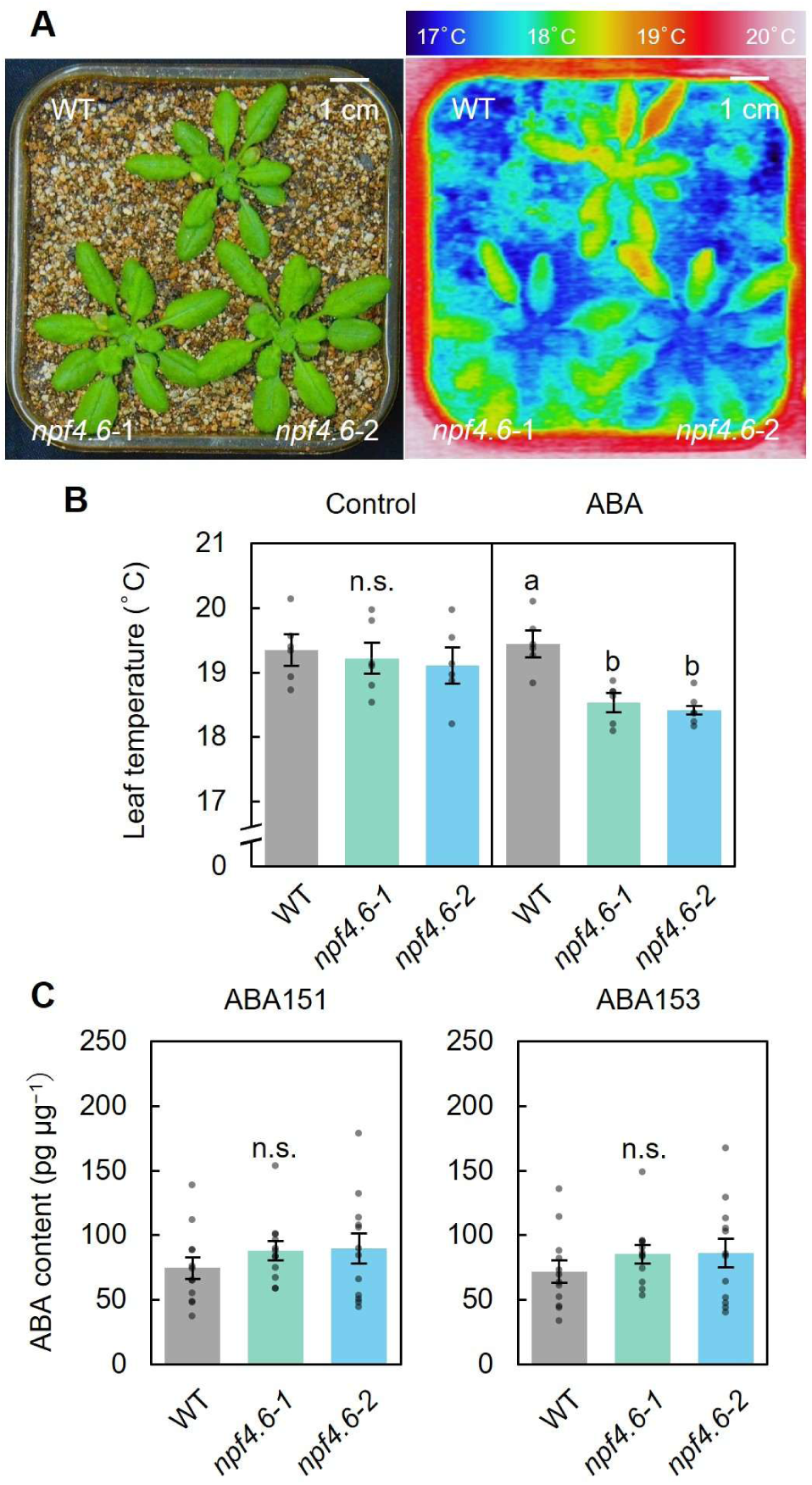
Evaluation of ABA transport by NPF4.6. (A) Representative plant growth and thermal image of WT and *npf4.6* mutants after ABA treatment and overnight incubation. Scale bars = 1 cm. (B) Leaf temperature of WT and *npf4.6* mutants (*n* = 6). (C) Quantification of ABA in xylem sap by LC-MS/MS (*n* = 10). ABA levels were calculated by monitoring either *m*/*z* 151.077 (ABA151) or *m*/*z* 153.092 (ABA153) as product ions derived from the precursor ion (*m*/*z* 263.1) of ABA. Values are mean ± SEM. Different letters indicate a significant difference at *P* < 0.05 by Tukey-Kramer HSD test.

### Comparison with other stomatal mutants

To assess the functional impact of NPF4.6 disruption relative to other well-established stomatal mutants, we compared leaf temperatures and drought tolerance among WT, *npf4.6-1*, *ost1*, and *aba3-1* (Fig. 5). Plants were well watered and grown for 4–5 weeks under PPFD of 130 μmol m^−2^ s^−1^, then exposed to PPFD of 500 μmol m^−2^ s^−1^ for 2 min before thermal imaging (Fig. 5A). Leaf temperatures did not differ between WT and *npf4.6-1*, whereas *ost1* and *aba3-1* showed significantly lower temperatures, indicating greater transpiration cooling by wider stomatal opening (Fig. 5B).

**Figure 5.**
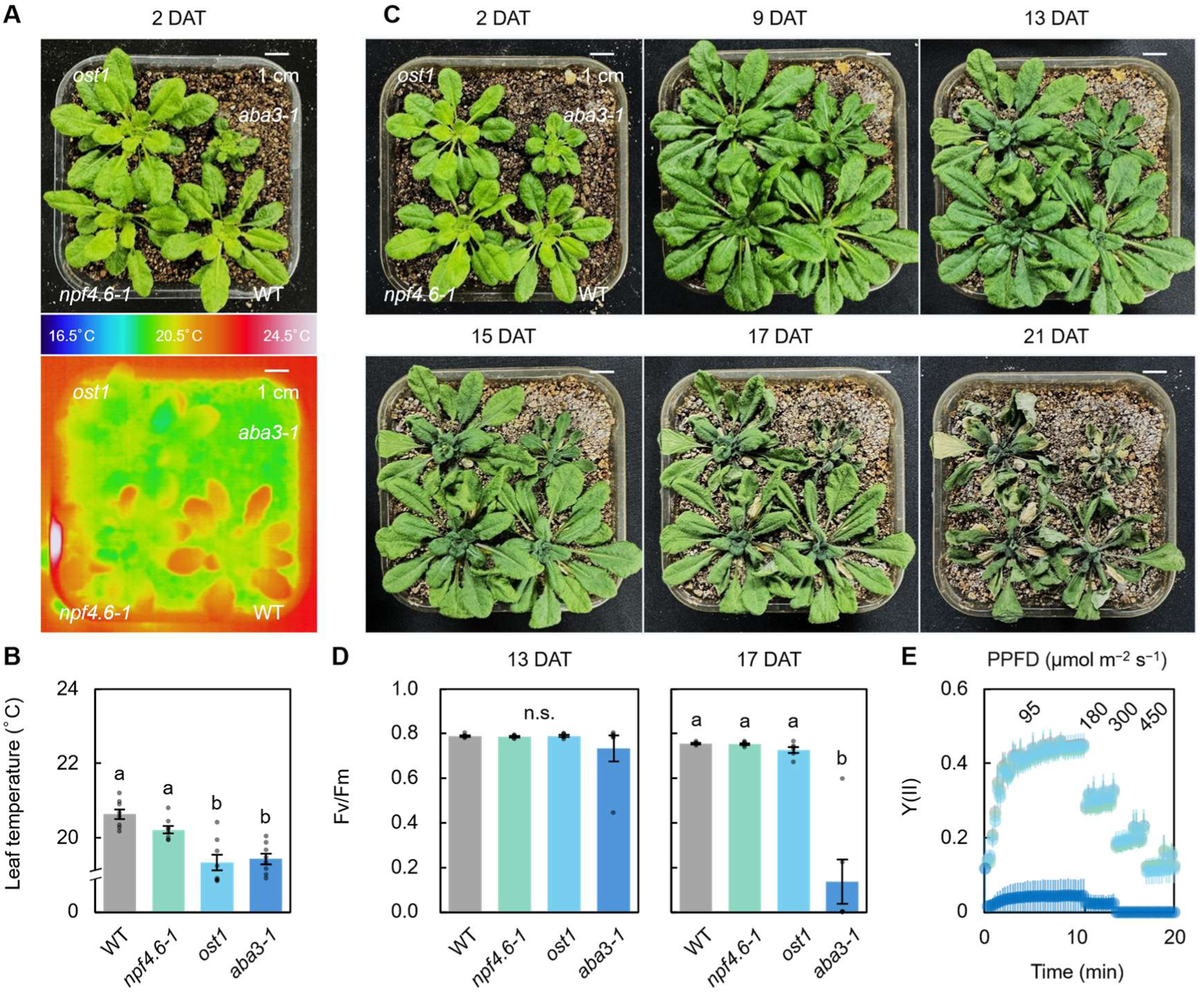
Transpiration and drought tolerance of *npf4.6* mutant and stomata-related mutants. (A) Representative growth and thermal image of WT, *npf4.6-1*, *ost1*, and *aba3-1* mutants under well-watered conditions (2 days after treatment, DAT). Scale bars = 1 cm. (B) Leaf temperature (*n* = 8). (C) Representative growth at 2, 9, 13, 15, 17, and 21 DAT. (D) Maximum quantum yield of PSII (Fv/Fm) at 13 and 17 DAT (*n* = 6). (E) Quantum yield of PSII (Y(II)) at 17 DAT (*n* = 6). Values are mean ± SEM. Different letters indicate a significant difference at *P* < 0.05 by Tukey-Kramer HSD test.

The same plants were then subjected to drought by withholding water for up to 21 days (Fig. 5C). *aba3-1* exhibited early wilting by 17 days after treatment (DAT) (Fig. 5C), and its maximum quantum yield of PSII (Fv/Fm) and quantum yield of PSII (Y(II)) was significantly lower than WT and other mutants (Fig. 5D, E). In contrast, *npf4.6-1* and *ost1* did not differ from WT in Fv/Fm and Y(II) under drought (Fig. 5C–E). These results indicate that the increase in *g*_sw_ in *npf4.6* mutants is moderate relative to strong ABA-deficient (e.g., *aba3-1*) or signaling mutants and does not compromise drought tolerance under the tested conditions.

## Discussion

In this study, we examined the role of the abscisic acid (ABA) transporter NPF4.6 in stomata-mediated gas exchange under FL by characterizing *Arabidopsis thaliana* loss-of-function *npf4.6* mutants. We found that NPF4.6 slows stomatal opening and CO_2_ assimilation after light irradiation, whereas *npf4.6* mutants exhibit faster photosynthetic induction and greater biomass under FL (Fig. 1, Fig. 2, Fig. 3). These results indicate that NPF4.6 adjusts *g*_sw_ in light-variable environments, revealing broader functions of ABA transport beyond drought responses.

### NPF4.6 modulates ABA-dependent stomatal kinetics under FL

In natural environments, light fluctuates rapidly due to clouds, canopy shading, and leaf movements (Pearcy, 1990; Slattery et al., 2018). Because stomata respond much more slowly than light changes, photosynthesis is often constrained (Kaiser et al., 2015; Tanaka et al., 2019; Lawson and Matthews, 2020; Tanigawa et al., 2024; Yoshiyama et al., 2024). Our results show that NPF4.6, an ABA importer (Kanno et al., 2012), regulates stomatal dynamics under these conditions. Specifically, *npf4.6* mutants showed faster stomatal opening and photosynthetic induction, and maintained higher *g*_sw_ under natural FL (Fig. 1, Fig. 3). A previous study reported that mutation of ABCG40, another ABA importer acting in guard cells, also leads to faster photosynthetic induction and increased shoot biomass under FL (Shimadzu et al., 2019). Together, these findings support a model in which NPF4.6 imports ABA into guard cells (Shimizu et al., 2021) to dampen rapid aperture changes during light transitions. Loss of this function is thought to reduce ABA levels in guard cells, accelerating stomatal opening and enhancing CO_2_ uptake. Future studies quantifying ABA in guard cells will clarify how NPF4.6 mediates this regulation.

### Distinct roles of ABA transporters under FL and drought

ABA is a central regulator of stomatal closure under soil water deficit, acting through enhanced biosynthesis in root or leaf vascular tissues and subsequently transported to guard cells (Kuromori et al., 2018; Hsu et al., 2021). Under severe drought, elevated bulk abscisic acid levels are thought to dominate stomatal regulation, potentially masking the contribution of individual ABA transporters because of redundant ABA transporters or other long-distance translocation mechanisms (Kanno et al., 2012; Hsu et al., 2021; Yang et al., 2024). Consistent with this view, *npf4.6* did not differ from the WT in growth or photosynthesis under drought conditions (Fig. 5), suggesting a limited contribution of NPF4.6 to drought tolerance under the tested conditions. By contrast, under well-watered conditions with FL, when bulk leaf ABA levels are low, spatial and temporal regulation becomes relatively more important for rapid stomatal adjustment (Bauer et al., 2013; Kuromori et al., 2018; Pridgeon and Hetherington, 2021). In our study, *npf4.6* mutants exhibited accelerated stomatal opening and maintained *g*_sw_ under FL (Fig. 1E–H), showing that NPF4.6-mediated ABA uptake into guard cells suppresses excessive aperture changes during light transitions. A similar phenomenon was observed in the *absg25* mutant, which displayed enhanced stomatal opening compared to WT under well-watered conditions but not under drought (Yang et al., 2024). Together, our results suggest that ABA transporters contribute differently depending on the environment: under severe drought, ABA biosynthesis dominates and transporter redundancy prevails, whereas under FL, when water is not limiting, NPF4.6 fine-tunes local hormone concentrations and response kinetics.

### Tissue-specific roles of NPF4.6 in ABA transport

*NPF4.6* is expressed in the vascular tissues of roots, hypocotyls, and leaves as well as in guard cells (Kanno et al., 2012; Shimizu et al., 2021). In this study, *npf4.6* mutants showed reduced stomatal closure compared to WT when exogenous ABA was applied from the root (Fig. 4A, B). However, ABA concentrations in the xylem sap collected from the stem base did not differ significantly between *npf4.6* mutants and WT (Fig. 4C). A previous study also reported that the movement of exogenously applied ABA in cut stems is comparable between WT and *npf4.6* mutants (Kanno et al., 2012). In addition, guard cell-specific complementation of *NPF4.6* partially restored low leaf temperature in the *aao3 npf4.6* double mutant, supporting a role of guard cell-localized NPF4.6 in facilitating ABA uptake and promoting stomatal closure (Shimizu et al., 2021).

Together, these observations suggest distinct tissue-specific functions among ABA transporters, with NPF4.6 likely to act primarily in guard cells as an ABA importer to regulate stomatal behavior, rather than influencing long-distance ABA transport via xylem sap.

### Environment-dependent trade-offs associated with NPF4.6 deficiency

The lower *iWUE* associated with higher *g*_sw_ in *npf4.6* can enhance carbon gain under FL when water is not limiting (Fig. 1), but may involve trade-offs depending on the severity and timing of drought stress during plant development (Leakey et al., 2019). From this perspective, retention of NPF4.6 may be advantageous for constraining stomatal opening under water limitation.

In addition to its role in ABA transport, NPF4.6 functions as a low-affinity nitrate transporter (Huang et al., 1999). A previous study reported reduced leaf area in *npf4.6* mutant compared to WT under high nitrate but not under low nitrate (Babst et al., 2019). In our conditions, *npf4.6* mutants accumulated more shoot and root biomass than WT (Fig. 2B, Fig. S1), but one line showed reduced leaf N concentration and N/C ratio (Fig. 2K, L), suggesting a partial impairment in nitrate uptake. Under high nitrate availability, the absence of NPF4.6 may reduce nitrate uptake, thereby offsetting the growth advantage of faster stomatal opening. Nitrate availability is also known to modulate ABA accumulation and plant responsiveness to ABA (Vidal et al., 2018; Ma et al., 2025). Although NPF4.6 itself does not transport nitrate and ABA competitively (Kanno et al., 2013), many ABA-responsive genes are induced under low nitrate but not high nitrate conditions, suggesting antagonism between nitrate and ABA signaling (Ma et al., 2025; Morales de Los Ríos et al., 2025). Thus, under high nitrate conditions, ABA responses may be suppressed, further limiting the effects of NPF4.6 deficiency on stomatal opening and growth.

Recent work also suggests that NPF4.6 exhibits low-pH-dependent and nitrate-promoted import activity for Na^+^, K^+^, and Li^+^ (Liu et al., 2025). Furthermore, overexpression of *NPF4.6* has been shown to enhance ABA-induced reductions in Cd, Mn, and Ni accumulation under heavy metal stress (Zhu et al., 2024). Therefore, future growth experiments at varying cation concentrations would help elucidate additional physiological roles of NPF4.6 and potential trade-offs associated with its loss of function.

### Agronomic relevance of NPF4.6 in field crop performance

Improving photosynthesis under dynamic light by accelerating stomatal responses is a key target for enhancing crop productivity (Slattery et al., 2018; Tanaka et al., 2019; Lawson and Matthews, 2020). Our results demonstrate that *npf4.6* mutants exhibit faster photosynthetic induction and greater biomass under FL conditions (Fig. 1, Fig. 2, Fig. 3), suggesting that modulating ABA transporters could enhance carbon assimilation in field-like environments. Unlike manipulation of ABA biosynthesis or major signal transduction components, which often results in severe dehydration and growth penalties (Léon-Kloosterziel et al., 1996), targeting transporters may allow finer control of hormone flux. Supporting this idea, the *A. thaliana* NPF4.6 homologue in tomato, SlNPF4.5 (Solyc05g006990), also functions as an ABA transporter, and its loss-of-function mutant exhibits reduced leaf temperature and higher germination rates under salinity stress, with only mild growth suppression (Shohat et al., 2020; Shohat et al., 2023). Moreover, NPF4.6 orthologs appear to be widely conserved in seed plants (Fig. S2). These results indicate that the function of NPF4.6 as an ABA transporter is likely conserved across species, making crop homologs potential targets for improving photosynthesis under field conditions. Future work should clarify how NPF4.6 expression and activity are regulated by natural environmental cues, and evaluate potential trade-offs associated with manipulating this pathway under agronomic conditions.

## Conclusion

This study identifies a role of the ABA transporter NPF4.6 in regulating stomatal dynamics in photosynthetic leaves under FL. Unlike during drought, where ABA biosynthesis predominates, light fluctuations require spatially regulated ABA transport to guard cells, where NPF4.6 plays a crucial role. Loss of NPF4.6 function accelerates stomatal opening, enhances photosynthetic induction, and increases biomass under dynamic light environments. These results highlight the environmental specificity of ABA transporter function and suggest that modifying hormone transport may improve photosynthesis and crop productivity in field-like conditions. Future studies should address the regulation of NPF4.6 and its functional conservation in crops.

## Materials and methods

### Plant materials and growth conditions

*Arabidopsis thaliana* wild type (WT, Col-0) and the *npf4.6* knockout mutants *npf4.6-1* (*ait1-1*) (SALK_146143) and *npf4.6-2* (*ait1-2*) (SALK_072696C), as well as *ost1* (SALK_008068), and *aba3-1* (CS157) were used. Plants were grown in plastic pots (280 mL or 380 mL) filled with a 1:1 (v/v) mixture of vermiculite and nutrient soil (Metromix 350; HYPONeX JAPAN, Japan). Growth conditions were 23°C, 65% relative humidity, 10-hour light period, photosynthetic photon flux density (PPFD) of 130 μmol m^−2^ s^−1^, and ambient CO_2_ concentration of 400 μmol mol^−1^. Pots were placed randomly on plastic trays and repositioned regularly. Plants were watered once a week.

### Gas exchange measurements

Photosynthetic induction was measured using plants kept in the dark overnight. Fully expanded young leaves from 6- to 8-week-old plants were used. Gas exchange was recorded every 30 s using a gas exchange system (LI-6400XT; Li-COR, USA) under the following conditions: air (block) temperature of 25°C, relative humidity of 55–65%, reference CO_2_ concentration of 400 μmol mol^−1^, and flow 500 μmol s^−1^. Leaves were kept in the dark for 5 min, then exposed to PPFD of 1000 μmol m^−2^ s^−1^ for 40 min.

Intrinsic water use efficiency (*iWUE*) was calculated as:

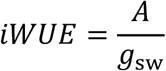

Normalized values for net CO_2_ assimilation rate (*A*) and stomatal conductance *(g*_sw_) were calculated as:

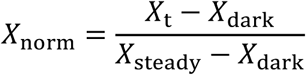

where *X*_dark_ is the 5-minute average under dark, *X*_steady_ is the 5-min average between 35–40 min under PPFD of 1000 μmol m^−2^ s^−1^, and *X*_t_ is the value at each time point. To calculate the photosynthesis induction, curve fitting was applied to the relationship between *X*_norm_ and time (*t*) using the following functions:

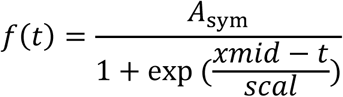

where *A*_sym_ is the maximum asymptote, *xmid* is the time at the inflection point, and *scal* is a scale parameter that determines the slope of the curve. Nonlinear least-squares fitting using the nls() function in R (v. 4.2.2). Optimal parameters (*A*_sym_, *xmid*, *scal*) were selected by minimizing the residual sum of squares between observed and predicted values. The following initial values were used: *A*_sym_ initial: 1.05 × the maximum of *X*_norm_; *xmid* initial: median of time values; *scal* initial: 1. The “port” algorithm was used for optimization, with a maximum of 10,000 iterations and a convergence tolerance of 1.0 × 10^−6^. Based on the fitted function, T_50_ and T_70_ were defined as the times required to reach 50% and 70% of *A*_sym_, respectively.

For measurements under simulated field light conditions, we used the light fluctuation pattern recorded in a rice paddy field in Kyoto Prefecture, Japan (35°02′N, 135°47′E) on 23 July 2016 (Kimura et al., 2020), which was converted to an 11-hour light period with a maximum PPFD of 500 μmol m^−2^ s^−1^. These fluctuations were reproduced in the LI-6400 cuvette (air temperature of 25°C, relative humidity of 55–65%, and reference CO_2_ concentration of 400 μmol mol^−1^), and photosynthetic parameters were recorded at 10 s intervals. Furthermore, normalized values for *A* and *g*_sw_ were calculated as:

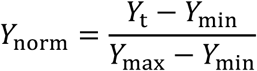

where, *Y*_min_ and *Y*_max_ are the minimum and maximum measured values of each parameter, and *Y*_t_ is the value at each time point.

### Growth analysis

Above- and below-ground tissues of 7- to 8-week-old plants were separated. Projected leaf area was measured using ImageJ (https://imagej.net/ij/). Dry weight was determined by drying samples at 80°C for 72 h. Leaf mass per area (LMA) was calculated as:

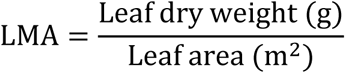

### Thermal imaging

Leaf temperatures of 5-to 6-week-old plants were measured using a far-infrared camera (Thermo Tracer TH9100MR; NEC Avio Infrared Technologies, Japan). For abscisic acid (ABA) treatment, a 10 μM ABA solution (A1049; Sigma-Aldrich Co. LLC, USA) was poured into the plastic tray the day before imaging, allowing the plants to absorb it from the bottom. Imaging was conducted the following day.

### Chlorophyll fluorescence measurements

Chlorophyll fluorescence was measured using a pulse-amplitude-modulated fluorometer (IMAG-MAX/L; Heinz Walz GmbH, Germany) in plants dark-adapted overnight. A saturation pulse was applied at an intensity of 10 for 720 ms, with measuring light intensity of 2, a frequency of 1 Hz, a gain of 1, and a damping of 2.

### Determination of nitrogen, chlorophyll, and Rubisco

Two leaf discs (8.5 mm in diameter) were obtained from a single fully expanded leaf of 7- to 8-week-old plants using a cork borer, frozen in liquid nitrogen, and stored at −80°C. Chlorophyll and Rubisco contents per unit leaf area were measured using methods described in previous studies (Porra et al., 1989; Kimura et al., 2020). For nitrogen concentration, three leaves dried at 80°C for 3 days were ground, and the nitrogen-to-carbon ratio in the sample was analyzed using an elemental analyzer (SUMIGRAPH NC-22F, Sumika Chemical Analysis Service, Japan) as described (Katsuhama et al., 2025).

### ABA contents analysis

The hypocotyls of 7- to 8-week-old plants were cut with a razor blade, and the detached shoots were inserted into 15 mL tubes. The tubes were centrifuged at 6800 ×g for 3 min at 5°C, and the liquid collected at the bottom of the tube was designated xylem sap. The collected xylem sap was then frozen at −80°C until the analysis. Ten microliters of xylem sap was mixed with 100 μL of d_6_-ABA (5 pg/μL in 50% (v/v) MeOH) (OlchemIm, Czech Republic) and 90 μL of MeOH, and filtered with Cosmo Spin Filter G (Nacalai Tesque, Japan) that had been pretreated with 200 μL of MeCN and then with 200 μL of 80% (v/v) MeCN. The filter was washed with 100 μL of 80% (v/v) MeCN, then combined with 200 μL of the flow-through from the previous step to yield a xylem sap sample. The sample was dried in vacuo and dissolved in 10 μL of water for LC-MS/MS analysis using a mass spectrometer (Q Exactive; ThermoFisher Scientific, USA) coupled to a UHPLC (Vanquish; ThermoFisher Scientific, USA). The samples were separated on an ACQUITY PREMIER HSS T3 column (1.8µm VanGuardFIT 2.1 × 100 mm; Waters, MA, USA) with a gradient of solvent B (0.01% formic acid in MeCN) in combination with solvent A (0.06% acetic acid), as indicated in Supplemental Table S1. The flow rate of LC was as indicated in Supplemental Table S2. The setting of mass spectrometer was as follows: mass tolerance (ppm) = 5, ion spray voltage (kV) = 2.5, collision energy (eV) = 10, S-lens radio frequency level = 50, resolution = 70,000, auto gain control target = 2 × 10^5^, maximum injection time (ms) = 200, isolation window (*m*/*z*) = 0.4, ion mode = negative. Data were processed with Xcalibur 4.4 (ThermoFisher Scientific, USA). d_6_-ABA was identified based on transitions of *m*/*z* 269.2 to 156.108 and 159.130. Endogenous ABA was identified based on transitions of *m*/*z* 263.1 to 151.077 and 153.092. Retention times of d_6_-ABA and ABA were 13.1 min. ABA levels were calculated based on the ratios of internal standard (d_6_-ABA) and non-labelled ABA.

### Statistical analysis

Statistical analysis was performed using R software (v.4.2.2; https://cran.r-project.org/bin/windows/base/). Data were analyzed by one-way ANOVA, followed by Tukey-Kramer HSD test (*P* < 0.05) or Dunnett’s test (*P* < 0.05).

## Acknowledgments

The authors thank Tomonori Niwa for his assistance with photosynthesis measurements.

## Data availability

All data are included in the manuscript and/or supporting information.

## Supplemental Figures

**Supplemental Figure S1.**
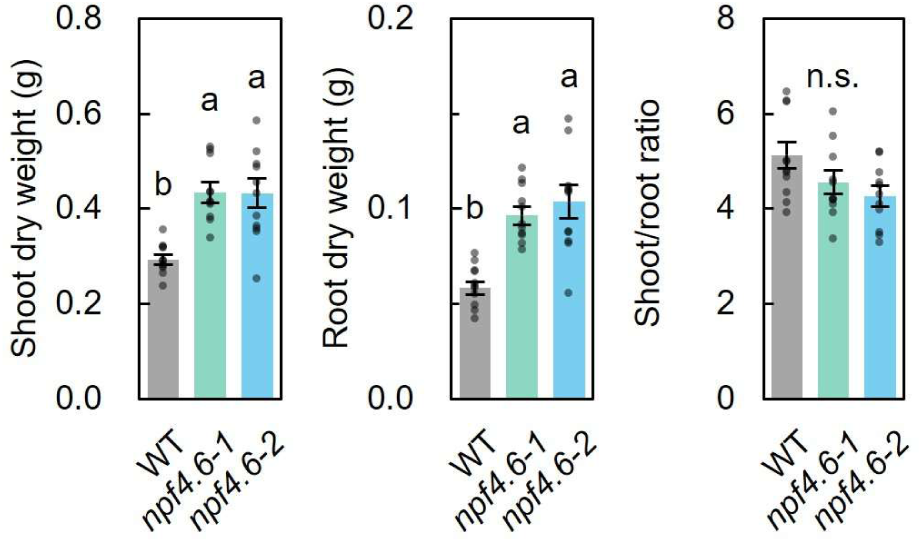
Growth of WT and *npf4.6* mutants under short-day conditions (second replication). Shoot and root dry weight was measured, and the shoot-to-root ratio was calculated in 7- to 8-week-old plants. Values are mean ± SEM. Different letters indicate a significant difference at *P* < 0.05 by Tukey-Kramer HSD test (*n* = 11).

**Supplemental Figure S2.**
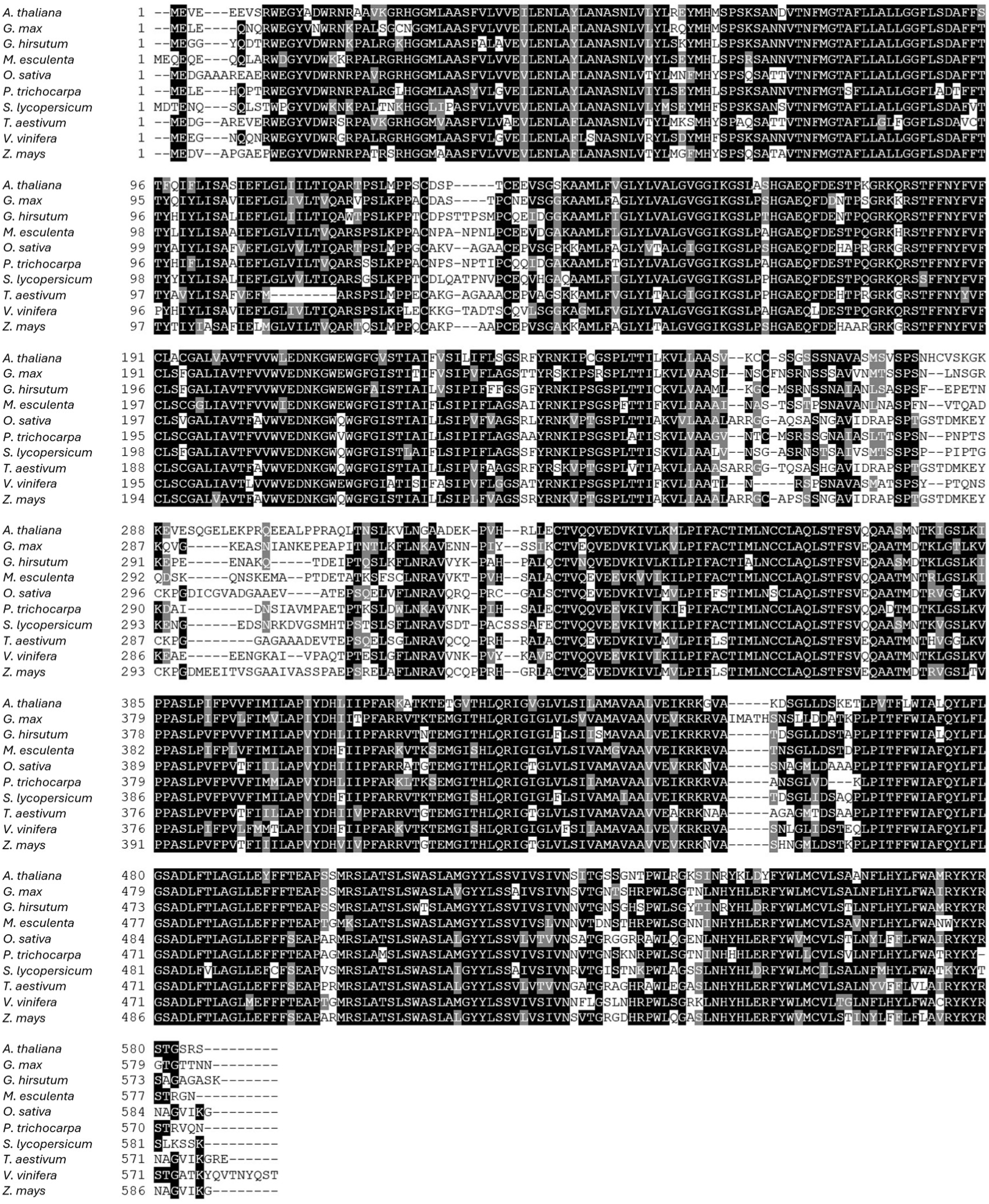
Multiple sequence alignment of *Arabidopsis thaliana* NPF4.6 and closely related proteins in crop species. Amino acid sequences of *Arabidopsis thaliana* NPF4.6 (NP_564978.1), *Glycine max* (XP_003551989.1), *Gossypium hirsutum* (XP_040940605.1), *Manihot esculenta* (XP_021613257.1), *Oryza sativa* (BAS98370.1), *Populus trichocarpa* (XP_006378161.1), *Solanum lycopersicum* (XP_004238765.1), *Triticum aestivum* (XP_044423362.1), *Vitis vinifera* (RVW82445.1), and *Zea mays* (NP_001150745.1) were aligned in MEGA12 (https://megasoftware.net/) using the MUSCLE algorithm. Conserved residues are highlighted in black, and similar residues are shaded in gray.

**Supplemental Table S1.**
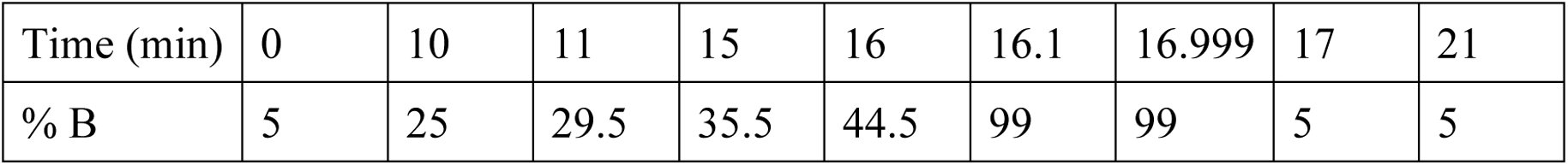
Percent of solvent B during the LC-MS/MS analysis.

**Supplemental Table S2.** Flow rate of LC during the LC-MS/MS analysis.

|  |  |  |  |  |  |  |
| --- | --- | --- | --- | --- | --- | --- |
| Time (min) | 0 | 16 | 16.1 | 16.9 | 16.999 | 21 |
| Flow rate (mL min<sup>-1</sup>) | 0.25 | 0.25 | 0.4 | 0.4 | 0.25 | 0.25 |

